# Greening conceals evergreening: contrasting trends for a socio-ecological system in Arctic Europe

**DOI:** 10.1101/2022.02.28.482210

**Authors:** M.W. Tuomi, T.Aa. Utsi, N.G. Yoccoz, C.W. Armstrong, V. Gonzalez, S.B. Hagen, I.S. Jónsdóttir, F.I. Pugnaire, K. Shea, D.A. Wardle, S. Zielosko, K.A. Bråthen

## Abstract

1. Ongoing Arctic greening can increase productivity and reindeer pasture quality in the tundra. However, greening may also entail proliferation of unpalatable species, with distinct consequences for pastoral socio-ecological systems (SES).
2. We show extensive greening across 20 reindeer districts in northern Norway between 2003 and 2020. The allelopathic, evergreen dwarf-shrub crowberry biomass increased by 60%, contrasted by smaller increases of deciduous dwarf-shrubs and stagnating forb and graminoid biomass.
3. We found evidence, although uncertain, of a negative relationship between biomass and reindeer densities, but only among forbs, the least abundant plant group.
4. Our results challenge the management decision-making, which aims at sustainable pasture management, but which assumes stationary density-dependent relationships. Changes in temporal vs. spatial relationships should be included as management criteria, to avoid mismanaging a SES in transition.
5. Large-scale shift towards increased allelopathy may undermine the resource base of a key Arctic herbivore and pastoral SES.

## Introduction

Rapid changes in the Arctic climate (IPCC 2021) are altering primary productivity, biodiversity and ecosystem functions (I. H. Myers-Smith et al. 2020; Barry, Berteaux, and Bültmann 2013; Post et al. 2019; Taylor et al. 2020), with effects cascading to and interacting with herbivore populations and coupled social-ecological systems (SES). Prevalent warming-induced vegetation trends include increased productivity, biomass and leaf area, identified as ecological greening of the Arctic (I. H. Myers-Smith et al. 2020), and shifts towards more resource acquisitive plant species (Bjorkman et al. 2018; Elina Kaarlejärvi, Eskelinen, and Olofsson 2013). In practice, such changes are expected to manifest as increased abundance of plants with taller stature and higher nitrogen (N) concentrations, especially willows and other deciduous shrubs, but also graminoids and forbs (Bråthen, Gonzalez, and Yoccoz 2018; Bjorkman et al. 2018; I. H. Myers-Smith et al. 2019). In areas where shrubs are already present or dominant, their growth can occur via increased cover, i.e. infilling of existing patches, or accumulation of biomass, i.e. vertical growth (I. H. a b Myers-Smith et al. 2011). Recent observations, however, indicate that Arctic vegetation changes conceal functionally contrasting trends. Field observations show increases especially in evergreen dwarf-shrubs across the circumpolar Arctic (Wilson and Nilsson 2009; Elmendorf et al. 2012; Maliniemi et al. 2018; Vowles et al. 2017; Vuorinen et al. 2017; Stewart et al. 2018). Evergreen plants often have high phenolic and low N content, and can contribute to lowering ecosystem productivity (Vowles and Björk 2019; Tuomi et al. 2018). An “evergreening” trend may therefore be functionally distinct from that of greening by deciduous plants, as it may suggest an ongoing decline in process rates, herbivore forage quality and biodiversity (Bråthen, Gonzalez, and Yoccoz 2018), despite increasing biomass in vegetation.

Proliferation of poorly palatable vs. palatable plants will likely have distinct consequences for herbivore populations and associated socio-ecological systems. In a rapidly changing Arctic, reindeer herding systems are readily subject to multiple climate and anthropogenic stressors that affect food availability (e.g. rain-on-snow events that lead to impenetrable ice layers that prevent foraging) and access to pasture (e.g. tourism and infrastructure) (Horstkotte et al. 2017; Hausner et al. 2020). Until now, however, changes in Arctic vegetation composition have been little considered among the novel threats to caribou and reindeer populations and indigenous pastoral systems (Post et al. 2009; Mallory and Boyce 2017; Horstkotte et al. 2017).

*Rangifer* (reindeer and caribou) are the most numerous Arctic ungulate, with grazing systems that span the circumpolar area (Mallory and Boyce 2017), and that are core to the indigenous Sámi livelihood and culture (James 2020). Akin to all extensive pastoral or grazing systems, *Rangifer* rely on high-quality plant resources (Sinclair and Krebs 2002; K. L. Parker, Barboza, and Gillingham 2009). Increasing plant productivity and abundance of N-rich and palatable herbaceous and deciduous species are expected to translate into higher survival and population growth rates of reindeer through positive bottom-up effects (K. L. Parker, Barboza, and Gillingham 2009; Tveraa et al. 2013). In contrast, poorly palatable species may reduce pasture productivity and quality to the extent that reindeer avoid areas where their dominance is high (Iversen et al. 2014). In a recent example, the satellite-derived greening signal in North-America was negatively related to caribou population growth rates, which was attributed to expansion of deciduous shrubs with high levels of anti-browsing defenses (Fauchald et al. 2017). The balance between the proliferation of poorly palatable and palatable plant species in *Rangifer* pastures across space and time, however, is poorly known.

Arctic terrestrial ecosystems often host strong herbivore-plant interactions, with co-occurring bottom-up and top-down dynamics (L. Oksanen and Oksanen 2000; Hoset et al. 2017; Tveraa et al. 2007). Arctic reindeer and caribou have highly context-dependent impacts on vegetation (Bernes et al. 2015), although they theoretically can strongly impact plant biomass along gradients of productivity through top-down effects (L. Oksanen and Oksanen 2000). In line with top-down effects, *Rangifer* have been found to modulate warming-induced vegetation changes. For instance, reindeer can counteract shrubification (Christie et al. 2015), e.g. keeping nutritious willows in a “browse-trap” (Bråthen et al. 2017), and thereby prevent the even more nutritious forbs and graminoids from being overgrown (Bret-Harte et al. 2001; Mod and Luoto 2016). Suppression of palatable species may, however, not be representative of how herbivores affect less palatable plants. For instance, the ability of *Rangifer* to modulate warming-induced changes in dominant, nutrient-poor evergreen dwarf-shrubs is less clear (Maliniemi et al. 2018; Vowles et al. 2017).

Central to many Arctic ecosystems and their change is the evergreen dwarf-shrub crowberry (*Empetrum nigrum*), a niche-constructing, allelopathic species (Bråthen, Gonzalez, and Yoccoz 2018; Wardle et al. 1998) (Supplementary Content Table). In Fennoscandia, crowberry is already abundant and commonly dominant. As a boreal-Arctic species it appears to thrive under warming climate in tundra and it tolerates various environmental stressors (Tybirk et al. 2000; Aerts 2010; Preece and Phoenix 2014; González et al. 2019). Owing to its low foliar nutrient concentrations, high levels of allelopathic polyphenolic compounds in its leaves and a dense, clonal growth form, crowberry is highly unpalatable, and can significantly retard ecosystem processes such as litter decomposition, soil nutrient fluxes and seedling establishment (Bråthen, Fodstad, and Gallet 2010; González et al. 2021; Tybirk et al. 2000; Wardle et al. 1998; Nilsson and Wardle 2005). Its dominance is related to suppressed biodiversity and herbaceous plant growth in summer pastures (Bråthen, Gonzalez, and Yoccoz 2018) (Supplementary Content Table), and reindeer have been found to avoid crowberry-dominated areas for most of the growing season (Iversen et al. 2014). Nevertheless, higher reindeer densities may be associated with spatially higher abundance of crowberry (Bråthen, Gonzalez, and Yoccoz 2018; Bråthen et al. 2007).

In the present work, we analyze vegetation across 20 reindeer districts in Norway to assess both temporal changes and spatial variation in reindeer summer pastures, and to what extent the associated Norwegian reindeer management (*Reindeer Husbandry Act* 2007) is capturing these changes (Fig. 1). The current reindeer management decision-making process aims at sustainable management of pastures (*Reindeer Husbandry Act* 2007; Ministry of Agriculture 2017, 32). However, the indicators used in the decision process do not measure the summer pastures and their quality directly, as the indicators are assumed to reflect the state of the system. Instead, the decision-making is based on an empirically supported negative density-dependent relationship between reindeer numbers and body mass as the focal indicator (Henden et al. 2020; Stien et al. 2021) (Fig. 1c, Supplementary Figure 2), and this relationship is assumed to be unaffected by long-term changes (Krebs 2002). Hence, the pastures themselves are not directly included in the decision-making process, as management indicators such as reindeer number and weights are easily accessible and they are linked to the economic valuation of the industry. The consequence is that long-term changes in productivity – a key determinant of sustainability of pastures and the husbandry – are managed through mandatory reductions of reindeer densities, easing the assumed top-down regulation of the pastures (Fig. 1b) whenever animal slaughter weights fall below a threshold. However, negative density-dependence in animal populations is not due only to changes in resources (Krebs 2002), and pasture productivity and composition may develop independently of animal densities, leading to bottom-up effects (Fig. 1b).

**Figure 1.**
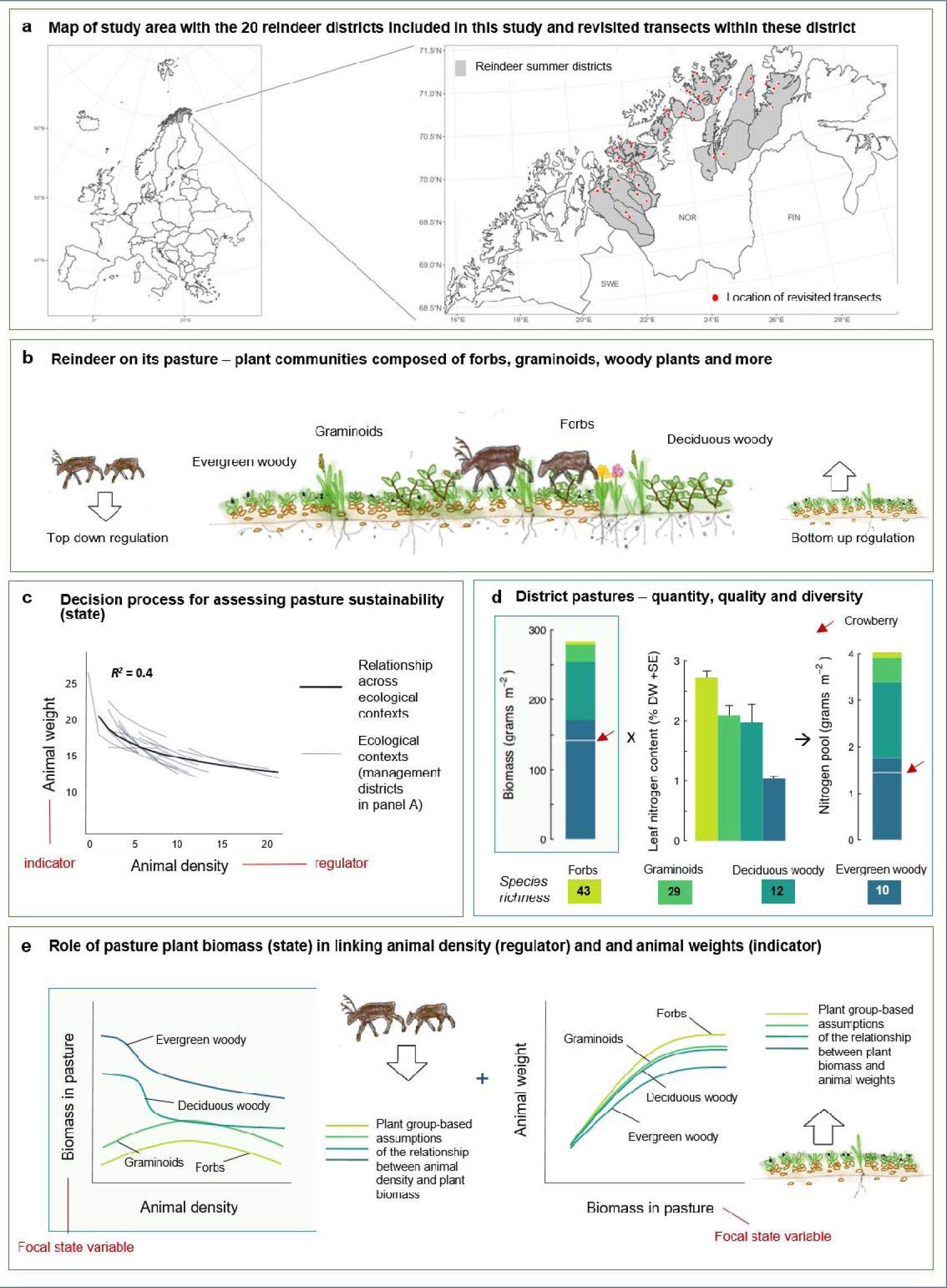
Role of pastures in the reindeer management system. The current management decision-making process of reindeer in Norway applies reindeer density as the regulatory mechanism of sustainable pastures (*Reindeer Husbandry Act* 2007). Autumn slaughter weight of animals, as a proxy for body condition, is used as the indicator for assessing sustainability of the summer pastures (state of the system), and no measures pertaining to the pastures themselves. **a)** The study area includes 20 management districts, spanning two latitudinal and six longitudinal degrees and representing different ecological contexts. Within each district we analysed transects for plant quantity, quality and diversity in 2003 and in 2020. **b)** Reindeer interact with their pasture through top-down and bottom-up effects. Approximate pasture functional group composition is visualized based on 2003 data. **c)** The management decision process applies animal weight as an indicator of pasture condition, here expressed as density-dependence of calf body mass in 2000-2019 within and across the management districts of the current study (Ministry of Agriculture, n.d.). **d)** In 2003 across the studied summer pastures, the average pasture consisted of plant functional groups of varying species richness (Bråthen et al 2007), nitrogen content (Murguzur et al 2019) and biomass (Bråthen et al 2007). Forbs, the most nutritious and species-rich group, made up the least biomass in the pastures. Conversely, the evergreen dwarf-shrubs, the least nutritious and species-poor group, made up the most biomass and with crowberry (red arrows) making up most of this biomass but not of the nitrogen pool. **e)** The role of plant biomass in linking reindeer densities and weights is indirect (*Reindeer Husbandry Act* 2007; Ministry of Agriculture 2017). The decision-making assumes a negative top-down relationship between animal density and plant biomass (left panel), and a positive bottom-up relationship between plant biomass and weights (right panel) (see Tveraa et al. 2013). Here, we inspect these assumptions further for plant functional groups. Left panel: Forbs and grasses, as more grazing tolerant groups, are expected to have a unimodal response to animal density (Wardle and Bardgett 2004). Deciduous shrubs are expected to show patterns of a browsing trap at densities above 5 animals/km ^2^ (Bråthen et al 2017), whereas evergreen shrubs are expected to be little affected from browsing (Iversen et al. 2014) but may decline under trampling effects (Tybirk et al. 2000). Right panel: Per biomass unit (right panel), we assume the most nutritious forbs to have the highest contribution to animal weights followed by graminoids, deciduous and evergreen shrubs in declining order of nutrient content respectively. Framed graphs in panels D and E are patterns and relationships that are empirically addressed here.

In contrast to the current management decision making, which emphasizes short-term and homogenous spatial and temporal effects, a community ecological perspective sees the pastures themselves as not uniform nor stable (Fig. 1b, d). There is a large variation in nutrient content between the plant functional groups in pastures (Fig. 1d), and the plant groups may respond differently to herbivory (Fig. 1e) and to other temporal and spatial factors, such as climate (temporal confounding) and bedrock nutrient content (spatial confounding). Therefore, plant responses to reindeer densities are likely to vary between plant functional groups in both space and/or time. Spatial and temporal patterns of plant groups and community compositions are therefore important determinants of changes in pasture quality (Fig. 1d), and hence of reindeer nutrition and growth (Fig. 1e). For instance, pastures abundant with the most nutritious plant groups could support high calf weights, whereas weights may saturate at lower levels if pastures consist mainly of crowberry (Fig 1d, e). In 2003, crowberry made up the majority of plant biomass across the northern Norwegian tundra, already resulting in pastures that were abundant in poor-quality forage of low nitrogen content (Fig. 1d).

We apply a unique, large-scale resurvey of 292 plant communities within 56 landscape areas sampled in 2003 and 2020 across 20 reindeer districts in northern Norway (Supplementary Figure 1) to assess variation in vegetation biomass relative to reindeer densities. Specifically, we ask 1) how preferred forage plants (forbs, graminoids and deciduous dwarf-shrubs) and less palatable evergreen dwarf-shrubs – with focus on crowberry – vary with reindeer density across space and in time, and 2) what are the implications of these changes for reindeer management decision-making. Given the ongoing warming trend in the region (Fig. 2), we hypothesize that all species and functional groups increase in abundance, including crowberry and other evergreen dwarf-shrubs. However, if the existing decision making process is adequate for achieving a sustainable management of pastures and the husbandry, observed changes in vegetation should be linked to reindeer density, or its changes. We thus ask if palatable plant groups have a more positive abundance response in management districts with low-to-intermediate or decreasing reindeer density (cf. Fig. 1e), and if evergreen dwarf shrubs have a more positive abundance response with high or increasing reindeer density. If the current management decision making is insufficient or incomplete, we expect opposite patterns (for dwarf-shrubs, Fig. 1e), or no association to reindeer density.

**Figure 2.**
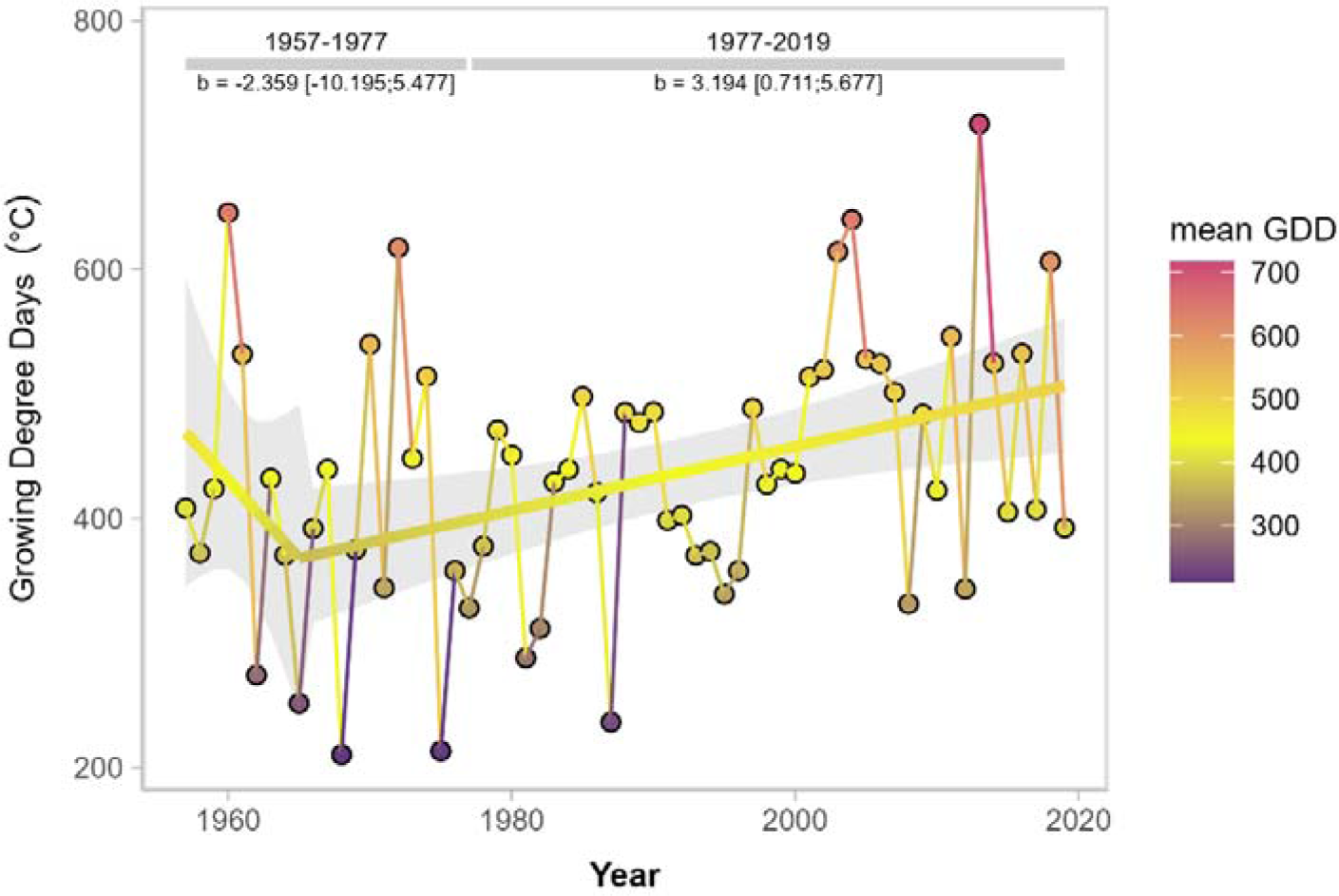
Growing degree days (GDD), as annual sums of daily temperatures above 5°C during May-October 1957-2019, averaged across the studied reindeer summer pastures (Pedersen et al. 2020/met.no). Solid thick line and shaded areas represent mean ± 95% confidence intervals from segmented linear regression with a breakpoint at year 1977. Parameter estimates are provided at the top of the figure.

## Methods

Located at the transition from subarctic to the low-Arctic, the Northern Fennoscandian study system is a bellwether for changes in tundra vegetation (Vuorinen et al. 2017; Elmendorf et al. 2012) and ecosystem functioning (T. C. Parker et al. 2021). The region has experienced lengthening of the growing season as spring and autumn have become warmer over the past century (Kivinen et al. 2017). With an approximate increase of 100 growing degree days in the studied summer pastures since the late 1970’s (Fig. 2), the pastures have been subject to a growing season prolongation, which is a key driver behind increased shrub growth in the Arctic (Post et al. 2019; Weijers et al. 2018; I. H. Myers-Smith et al. 2019). We revisited a hierarchical sampling design from 2003, with plant communities nested in blocks and reindeer districts (Bråthen et al. 2007) (Supplementary Figure 1). We used identical georeferenced plant communities and sampling methods within each community in both 2003 and 2020. Each plant community was represented by a 50m long transect with 11 equidistant, triangular measurement plots with 40cm sides (Bråthen et al. 2007) (Supplementary Figure 1). We analyzed plant data as biomass (g/m ^2^) and cover (number of plots within transect with observed presence).

We decomposed the reindeer density to its spatial, temporal, and residual components (Oedekoven et al. 2017), and standardized all three predictors to a mean of 0 and variance of 1 for better effect comparability and model convergence. The temporal component represents the change in average reindeer densities between the two time points, the spatial component the difference in the time-averaged district reindeer densities and the residuals the district-specific change in reindeer densities that is not captured in the average spatial and temporal densities. Models for forbs and graminoids included a quadratic term for the spatial component, while deciduous dwarf-shrubs and crowberry included a linear term only (cf. Fig 1e), and all models included block and district as group-level intercepts. We fitted Bayesian generalized linear mixed models (Bürkner 2017) with weakly informative default priors, and checked model convergence and independence of HMC chains based on the 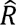 statistics (< 1.002) and effective sample size (> 2000) (Muth, Oravecz, and Gabry 2018). Lack of spatial autocorrelation in group-level effects and model residuals was assessed visually. We used averaged reindeer densities from the previous 23 years for 2003 (1980-2003, same data as in Bråthen et al. 2007) and 17 years for 2020 to avoid temporal overlap in the data (2003-2019, Ministry of Agriculture, n.d., accessed via reinbase.no).

## Results

Across the entire study area, crowberry standing biomass increased by 60% (Fig. 3a, Supplementary Result Table 1A), and cover by 14% (Supplementary Result Table 1B and 2) from 2003 to 2020. The increase of deciduous dwarf-shrub biomass was nearly an order of magnitude smaller than that of crowberry’s (Fig. 3a, Table 1), and we found no change in the cover of deciduous dwarf-shrubs. The average reindeer densities declined from 6.47 to 6.24 animals/km ^2^ over the same time period. The change in crowberry and deciduous dwarf-shrubs and shrubs was spatially consistent, as they increased in biomass in 90% and 85% of the districts, respectively. The increase in the woody shrubs’ biomass, crowberry in particular, was in sharp contrast with no temporal effects on the biomass and cover of forbs and graminoids (Fig. 3a, Table 1, Supplementary Result Table 2). The frequency of pasture communities in which more than 25% of total vascular biomass was crowberry rose over this period from an already high 0.76 to 0.83.

**Figure 3.**
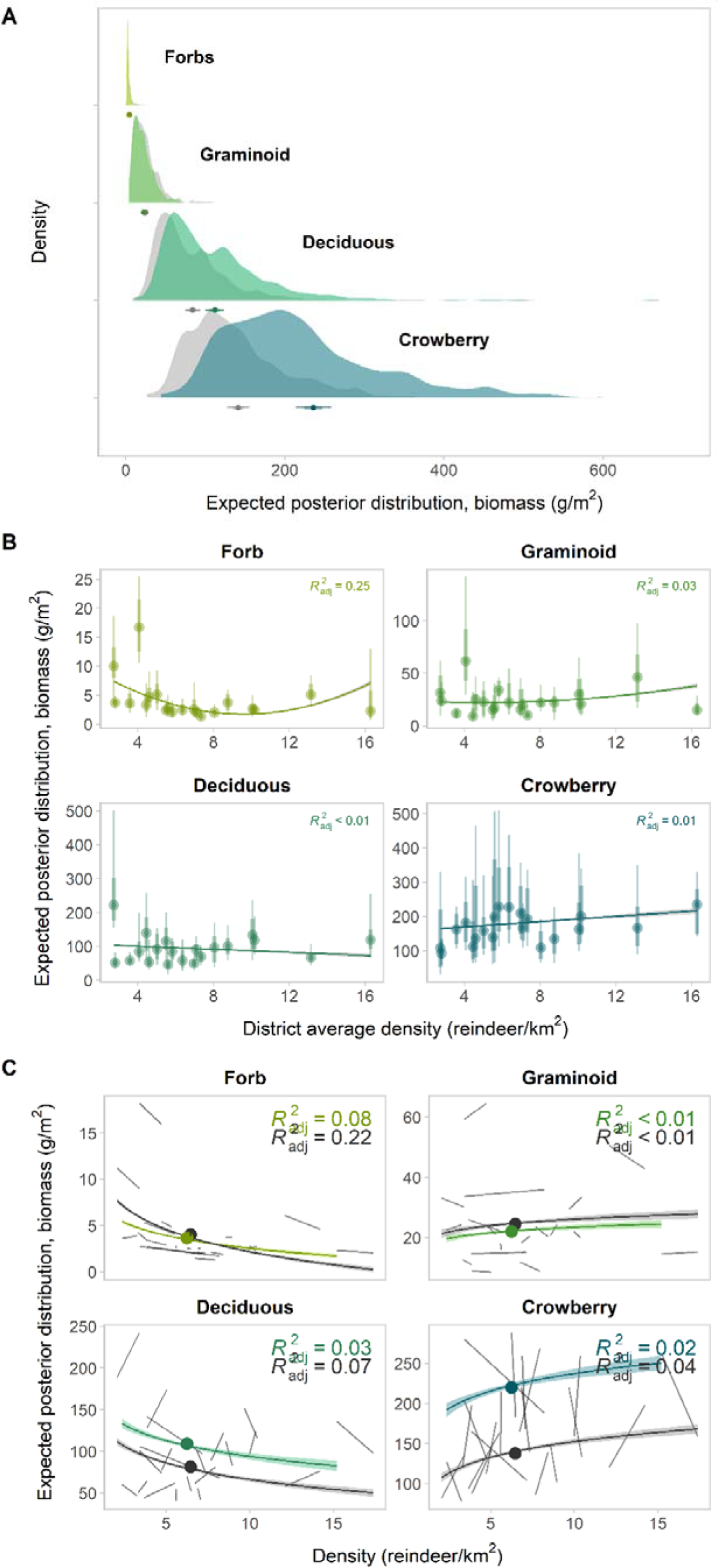
Modeled temporal and spatial variation in plant group biomass (forbs, graminoids, deciduous shrubs and dwarf-shrubs and crowberry). **A)** Estimates of posterior mean distributions of biomass of forbs, graminoids, deciduous woody plants and crowberry, separately for each plant group in 2003 (grey) and in 2020 (greens). Means and 95% confidence intervals from observed data for each year are provided as point intervals below each biomass density plot (note full overlap for forbs and graminoids). **B)** Association between plant group biomass (mean and model-derived 95% and 80% credible intervals of the estimated posterior mean distribution) and the district-averaged reindeer density. Each point represents a reindeer district. **C)** Model-derived estimates of the relationship of plant biomass and the (recomposed) reindeer density. Each short solid black line connects averaged model estimates in 2003 and 2020 in each district, and a regression line is fitted with log(density) across all districts. In panels B and C, the the coloured regression lines and adjusted R^2^ across all districts are estimated with the package *ggpmisc* functions stat_poly_line and stat_poly_eq (Aphalo 2023).

**Table 1.** Model parameter estimates and credible intervals, with plant functional group biomass as response and decomposed reindeer density as predictors. Note that the temporal coefficient links with decrease in reindeer density over time, and hence a negative value indicates an increase over time. For estimates of group-level standard deviation (random effects), see Supplementary Results Table 3. Error terms Q2.5 and Q97.5 represent the 95% credible interval. Bold font indicates strong support for the effect, and italic indicates relatively strong support for the effect, given the data.

| Response | Parameter | Estimate | Q2.5 | Q97.5 |
| --- | --- | --- | --- | --- |
| Biomass ~ spatial + spatial<sup>2</sup> + temporal + residual |  |  |  |  |
| Forbs | Intercept | 1.777 | 1.394 | 2.168 |
|  | *Spatial* | -0.424 | -0.831 | -0.006 |
|  | Spatial<sup>2</sup> | 0.205 | -0.071 | 0.475 |
|  | Temporal | 0.041 | -0.093 | 0.171 |
|  | Residual | -0.069 | -0.161 | 0.020 |
| Graminoids | Intercept | 3.132 | 2.799 | 3.471 |
|  | *Spatial* | 0.012 | -0.378 | 0.399 |
|  | Spatial<sup>2</sup> | 0.037 | -0.213 | 0.286 |
|  | Temporal | 0.041 | -0.072 | 0.154 |
|  | Residual | -0.013 | -0.088 | 0.061 |
| Biomass ~ spatial + temporal + residual |  |  |  |  |
| Deciduous woody | Intercept | 4.491 | 4.251 | 4.791 |
|  | *Spatial* | 0.028 | -0.224 | 0.280 |
|  | **Temporal** | **-0.220** | **-0.302** | **-0.137** |
|  | Residual | -0.028 | -0.082 | 0.027 |
| Crowberry | Intercept | 5.229 | 5.079 | 5.399 |
|  | *Spatial* | 0.081 | -0.092 | 0.254 |
|  | **Temporal** | **-0.340** | **-0.420** | **-0.261** |
|  | Residual | 0.001 | -0.052 | 0.054 |

While the overall temporal change in reindeer densities did not translate to change in forbs, we found that the spatial pattern in district-level reindeer densities had a negative relationship with forb biomass (Table 1). This effect had, however, a relatively high uncertainty (Table 1), and it tended to become weaker in 2020 (Fig. 3c). The shape of the relationship was also different from the expectation, with lowest estimated biomass at intermediate reindeer densities and a non-significant quadratic term (Table 1). There was no evidence that the spatial differences in reindeer densities explained variation in any of the other plant groups.

In the studied reindeer districts over the study period, densities declined in 50%, increased in 45% and stayed stable in 5% of reindeer districts (calculated with a change threshold of 0.1 reindeer/km ^2^). None of these district-specific changes in density explained variation in any of the plant groups (Table 1, Supplementary Figure 3). In our dataset, the spatial scale for important group-level effects differed between plant groups. Crowberry biomass varied substantially among landscape areas within reindeer districts, but little at the scale of districts (Fig. 3b, Supplementary Result Table 3). Deciduous shrubs and graminoids varied markedly between both districts and among landscape areas. In contrast, variation in forb biomass was largest between – and not within – districts (Fig. 3b, Supplementary Result Table 3).

## Discussion

In line with our first expectation, we found substantial increases in woody vegetation biomass. The difference in magnitude between the evergreen and deciduous woody plant proliferation, however, was surprising, with pasture evergreening far outpacing greening by deciduous shrubs. We found no evidence that variation in woody species linked with spatial density patterns nor with district-specific changes in reindeer densities. Therefore, it is plausible that the observed temporal changes are not attributable to the reindeer density change *per se*, but are mainly confounded by other temporal effects, such as changes in climate (Fig 2), corroborating the hypothesized evergreening and greening trends. Proliferation of crowberry appears to mainly occur through accumulation of biomass (vertical growth), but also through infilling, whereby ever larger surface areas are potentially impacted by crowberry.

In contrast to the expectation of temporal change, the least abundant, yet most species-rich and productive plant groups, the forbs and the graminoids, show no change over 18 years. This result was also surprising, given the overall decrease in reindeer densities and an ongoing warming trend, which both could be expected to favor herbaceous growth forms. Our results indicate that the spatial negative relationship of forbs and reindeer density may not be informative to predict changes over time. We note that the abundance of forbs and graminoids is extremely low, their rarity making assessment of patterns difficult at a large spatial scale and increasing uncertainty of the estimates. Nevertheless, the persisting rarity of the most productive plant groups combined with rapid proliferation of the least palatable evergreen dwarf-shrubs across landscapes is likely of high importance for pasture management and the pastoral SES.

Our results point at evergreening and crowberry proliferation as a potential major bottom-up forcing on the pastures (Fig. 3a, c) (cf. le Roux et al. 2013), effectively decoupled from district-level variation in reindeer density. Sustainable management of tundra pastures and the pastoral SES will be contingent on models that are representative of the managed system and its spatial and temporal uncertainties (Milner-Gulland and Shea 2017; Regan, Colyvan, and Burgman 2002; Latombe et al. 2019). An insufficient decision making process that misses influential variables or processes may severely undermine management objectives (Madden and McQuinn 2014), and lead to non-resilient management efforts, reducing the capacity of the system to adapt and maintain system resilience against undesirable states (Dudney et al. 2018). A pastoral management practice that does not monitor changes in the plant resource *per se*, such as the Norwegian reindeer management decision-making process, would likely remain functional or even thrive under increasing productivity and greening of palatable plants. However, managers would be ill equipped to detect and manage bottom-up effects that slowly reduce pasture quality. In the following, we argue that if left unchecked, allelopathic evergreening may have severe, adverse, long-term consequences for the diversity, productivity and resilience of reindeer pastures and the pastoral SES.

### Evergreening: allelopathy and the mechanisms of transition

As a niche-constructing species, crowberry can modify the environment once established (González et al. 2021; Bråthen, Gonzalez, and Yoccoz 2018; Wardle et al. 1998), as allelochemicals in crowberry’s leaves and accumulating litter can push the system towards a state of strong allelopathy (Fig. 4). The growth-inhibiting effects of crowberry litter can remain after the plant itself is gone (Pilsbacher et al. 2021; Dorrepaal, Cornelissen, and Aerts 2007; Aerts 2010), giving rise to legacy effects. Diminishing diversity due to allelopathy happens gradually through reduced seedling recruitment (González et al. 2015; 2021), and over longer time scales induce an extinction debt on the local plant communities (Kuussaari et al. 2009; Gilbert and Levine 2013), as local seedbanks disappear (Fig. 4a). These legacy effects, along with the potential longevity, recovery potential, dense growth and poorly palatable leaves of crowberry (Aerts 2010), suggests crowberry dominance of communities is likely to be a highly resilient state (E.a Kaarlejärvi, Hoset, and Olofsson 2015; González et al. 2021; Wardle et al. 1998; Nilsson and Wardle 2005). Once established, the state may require strong external disturbances to reverse (Fig. 4b, left panel). Consequently, we hypothesize that the long-term effects from crowberry allelopathy at the community- and ecosystem-level may represent an ongoing shift with potential context-specific thresholds (Hughes et al. 2013) (Fig. 4b, left panel).

**Figure 4.**
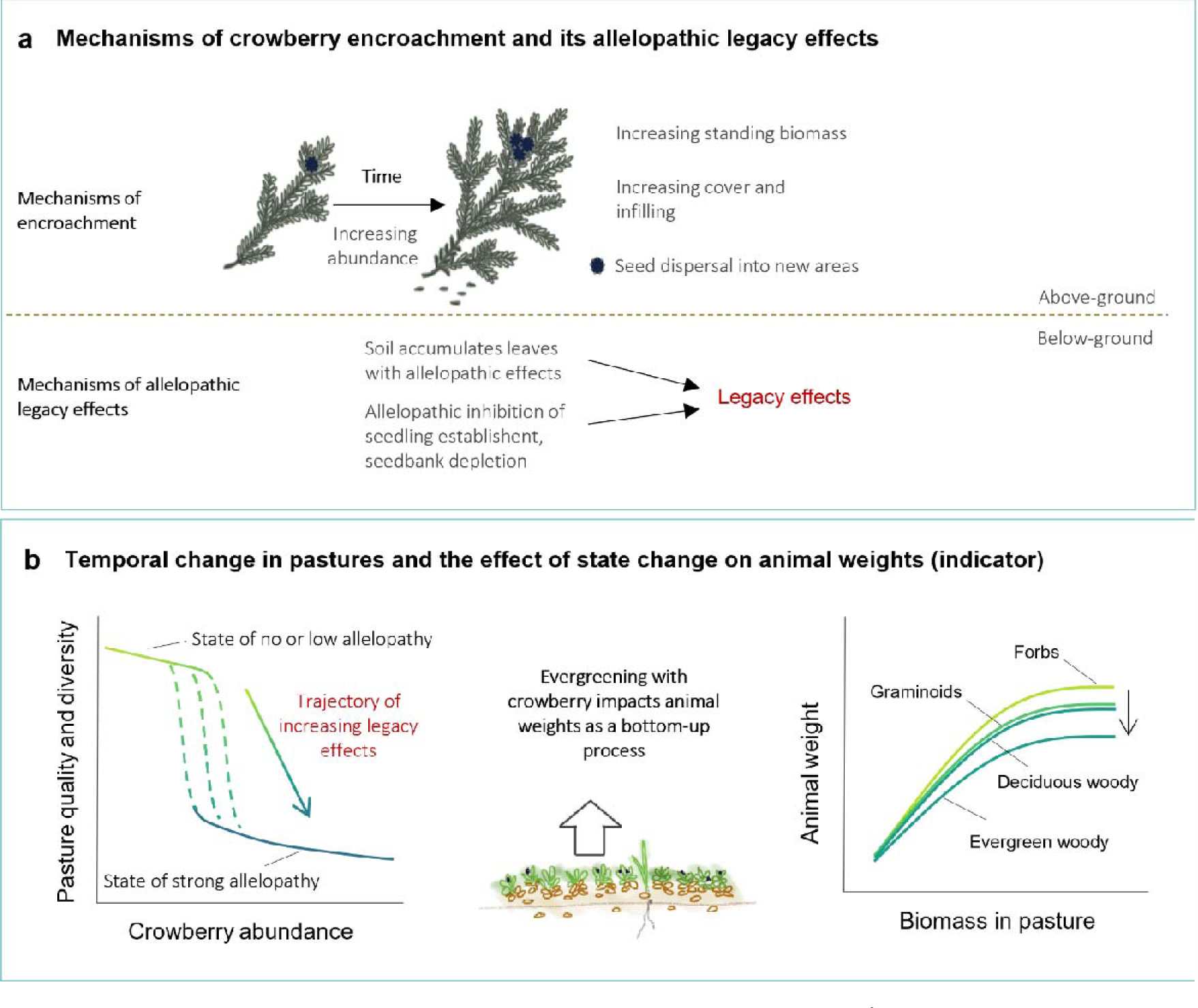
Mechanisms and consequences of crowberry proliferation. **A)** Mechanisms of crowberry proliferation and increased allelopathy over time. Already an abundant and long-lived species, slow growth may still cause substantial increments in abundance. Growth of established plants results in infilling and increased biomass, leading to increasing allelopathic litter effects in the soil. In addition, dispersal via seeds and clonal reproduction are means of lateral encroachment, increasing the areal extent of the plant. **B)** Potential transition in pasture state and its implication for the state-indicator relationship. Left panel: Increasing crowberry abundance may push the system towards a state of strong allelopathy. The state may be highly resilient, with litter and seed bank -mediated legacy effects that over time will reduce new plant establishment (Gonzalez et al 2021) at levels as low as 25% crowberry out of total community standing biomass (Bråthen et al 2018). Right panel: Conceptual model of effects of evergreening and crowberry proliferation on reindeer weights. Relationship between pasture biomass and calf weights is dependent on plant composition, and unlikely to be invariable over time. Changing bottom-up effects and transition to increasing allelopathy in pastures may have adverse, yet slow impacts on reindeer condition.

While we show that crowberry proliferation occurs in landscapes across northern Fennoscandia, we also find spatial variability in its increase especially at relatively local landscape scales, indicating there are ecological contexts in which the shift is stronger, weaker, or absent. Tundra vegetation changes are repeatedly found to be spatially heterogeneous across scales (Bjorkman et al. 2018; Van Wijk et al. 2004; Graae et al. 2018) and can be linked to e.g., variation in microclimate or herbivory. For instance, cyclic small rodent outbreaks across the resurveyed region can decimate dwarf-shrubs and especially crowberry in patches across landscapes (Tuomi et al. 2018; Hoset et al. 2017) (Supplementary Figure 4) - a temporary reduction even detectable from space (Olofsson, Tømmervik, and Callaghan 2012). However, strong localized small rodent grazing has not limited the overall high-magnitude, long-term encroachment of crowberry documented here.

While neither small nor large herbivores may halt the overall trend of crowberry encroachment across tundra pastures, they are also likely adversely affected by the observed low abundance of forbs and diminishing prevalence of communities with little crowberry (T. Oksanen et al. 2020; Iversen et al. 2014). Apart from reindeer, other endotherm herbivores, such as small rodents, ptarmigans, domesticated sheep and musk-ox also rely on N-rich forage (Schmidt et al. 2018), and seek pastures with high-quality food (Iversen et al. 2014; Jenkins et al. 2020; Skarin et al. 2020). Furthermore, crowberry being a wind-pollinated plant (Bell and Tallis 1973), its encroachment may over time also affect insect pollinators. In summary, crowberry encroachment of the magnitude documented here – and predicted e.g. for Arctic Greenland (Stewart et al. 2018) – raises significant concerns of cascading effects onto the tundra biota, ecosystem and human beneficiaries relying on them, including pastoralists, sheep farmers and game hunters.

### Way forward: monitor and manage

In the context of the reindeer management system, an evergreening trajectory towards higher evergreen dwarf-shrub (particularly crowberry) abundance with potentially lowered pasture process rates, forage availability and carrying capacity, threaten a risk of mismanagement and loss of resilience of the pastoral SES (Valéry, Fritz, and Lefeuvre 2013; Carey et al. 2012; Shackleton, Shackleton, and Kull 2019) (Fig. 4b, right panel). Our results suggest that revision of the current reindeer management decision-making process should add an adaptive component sensitive to long-term, decadal changes to the current approach focusing on relatively short-term fluctuations. Inclusion of measures relating to pasture plant diversity and productivity, in line with intentions for sustainable reindeer husbandry (*Reindeer Husbandry Act* 2007) and for translating the IPBES Global Assessment to policy (Ruckelshaus et al. 2020), will be crucial. In this regard, particular attention should be given to the most species rich and nutritious, yet scarce growth form, the forbs (Bråthen, Pugnaire, and Bardgett 2021).

Presently, however, crowberry proliferation, and its ecological and socio-economic impacts, remain poorly understood; this can impede development of novel management norms and objectives (Vaas et al. 2021; Davis et al. 2019). First, general acceptance and evidence-based baselines are often difficult to establish for slow and poorly detectable changes (Schneider, Leifeld, and Malang 2013), such as the creeping infilling and biomass accumulation of a slow-growing dwarf-shrub. The trajectory of evergreening is likely long, but large-scale empirical evidence goes back barely half-a century (Maliniemi et al. 2018). Second, contemporary validation and monitoring of Arctic vegetation change has relied heavily on remote sensing indices (I. H. Myers-Smith et al. 2020). Remote sensing may not necessarily capture changes in functional composition (Wang et al. 2021), distinguish greening and evergreening as functionally different processes (Vowles and Björk 2019), or document the species diversity of pastures. Adaptive and long-term ecosystem and pasture monitoring programmes are key to mitigating uncertainty about climate-driven vegetation change (Post et al. 2009; I. H. Myers-Smith et al. 2019; Lindenmayer and Likens 2009). Such programmes have recently been implemented, for example, in Iceland (Marteinsdóttir et al. 2021), and in the Varanger peninsula and Svalbard, in Norway (Ims and Yoccoz 2017).

Our results suggest that the current regulator of the management system (reindeer numbers) may not function in an efficient manner to support resilient and sustainable pastures. Additional management strategies, interventions and indicators that directly address spatial and temporal variation in diversity, productivity and heterogeneity of pastoral landscapes are needed. Such strategies have centennial or even millennial roots in extensive management practices of European coastal heathlands (Webb 2008; Vandvik et al. 2005) and in European agri-environmental policy that has focused on preventing loss of open semi-natural grasslands (Critchley, Burke, and Stevens 2004). Fire has the capacity to ameliorate soil conditions against crowberry’s allelopathic effects (Bråthen, Fodstad, and Gallet 2010; Keech, Carcaillet, and Nilsson 2005), suggesting management through burning as one potential way to locally control ecosystem state shifts similar to those in coastal heathlands (Vandvik et al. 2005) or boreal *Pinus*-dominated forests (Wardle et al. 1998; Nilsson and Wardle 2005). Here, the challenge lies in the already scarce and diminishing herbaceous resource, the forbs (Bråthen, Pugnaire, and Bardgett 2021). Scarcity begets scarcity through increasing seed limitation under encroaching crowberry dominance and allelopathic effects (González et al. 2021), and promotion of productive vegetation would likely require re-building seed banks alongside soil amelioration. Spatial rarity also poses challenges for monitoring efforts. Despite such challenges, we believe a resilient management approach to tundra reindeer pastures and pastoral SES is both attainable and urgent.

### Conclusions

Evergreening puts an increasing pressure on sustainable land-use planning and prioritization to preserve remaining forb and graminoid-rich pastures (Webber et al. 2022) and the biodiversity of tundra landscapes against rapid homogenization (Stewart et al. 2018). Evergreening through crowberry encroachment exemplifies emergence of super-dominance among native species, a phenomenon linked with anthropogenic pressures or novel climates across biomes (Pivello et al. 2018; Shackelford et al. 2013; Valéry et al. 2009), with ecological consequences not unlike those of non-native invasive species.

Overall, our results on crowberry encroachment suggest an ongoing decline in the quality of pasture land (Skarin et al. 2020; Iversen et al. 2014), adding to ongoing loss of land with productive pastures due to human disturbance (Hausner et al. 2020) and other stressors (Vaas et al. 2021; Horstkotte et al. 2017). Loss of pasture quality may additionally amplify other stressors, further eroding the resilience of an increasingly vulnerable Arctic system.

Ignoring the capacity of native species to be drivers of change is a critical blind spot that threatens the biodiversity and sustainability of ecosystems. For the Arctic pastoral system studied here, the management of tundra ecosystems and pastoral SES post-2020 should align targets of reindeer and biodiversity management and ensure resilience through ecosystem-based monitoring of the biodiversity of pastures. Only with such comprehensive approaches will sustainable management of diversity in complex social-ecological systems be possible in changing climates.

## Methods

### Study area and resurvey design

We conducted a vegetation survey of 292 remote, georeferenced vegetation communities in summer pastures across Northern Norway during peak growing season in 2003 and again in 2020. The study area (lat N69° 25.806’-N70° 58.471’, lon E20° 47.186’-E27° 31.099’) includes strong climatic gradients from west to east and from coast to inland, altitudinal variation from 60 to 600 m asl, as well as variation in bedrock types. Sampling included the most common vegetation types in the region: heaths, mires, snow beds, meadows, and windblown ridges. The survey design (Supplementary Figure 1) incorporates summer pasture areas of 20 reindeer districts, where adjacent districts of similar climatic conditions were organized into 10 district pairs of contrasting reindeer densities, based on average reindeer density from 1980 to 2003. Within each district, a 2×2 km vegetated grid was assigned out of which a random subset was chosen as landscape areas for sampling. Within each landscape area, a random set of 25 of the 100, 200 x 200m squares that constitute the vegetated grid were assigned for plant community sampling. Squares were then sampled by the transect method (each transect representing a plant community), whereby a 50m transect was placed from the midpoint towards a random GPS-position along a circle with a 50m radius. We used the point-frequency method in 11 plots every 5m along the transect to sample each community (Supplementary Figure 1). Each plot was measured by placing a triangular frame with sides of 40cm and one pin in each corner, counting all intercepts with the vegetation (Bråthen and Hagberg 2004). The a priori stratification of the original 2003 design, inclusion rules of transects and sampling method are described in more detail in Bråthen et al. (2007, 2018). In 2020, we resurveyed a subset of 292 (of the original 1450) communities in 56 (of if the original 151) landscape areas. The re-sampling retained the geographic extent as well as most of climatic and abiotic variability. Based on transect descriptions from 2003 (e.g., “transect was moved 10m backwards due to a lake”) we deemed the relocation of communities accurate.

### Environmental data

We estimated climatic trends between 1957 and 2019 in the studied summer pastures for growing degree days (GDD), using data described in Pedersen (2021). We applied segmented linear regression models using the R-package segmented (Muggeo 2017) to explore trends and breakpoints in the mean estimates for the regional climate.

We retrieved data on reindeer numbers from the onset of the reindeer herding year (cf. Stien et al. 2021) for each studied reindeer herding district (Ministry of Agriculture, n.d.), which we then divided by summer pasture area (km ^2^) to obtain reindeer density for each district (individuals/km ^2^). We used the average density from 1980 to 2003 as a predictor for the 2003 vegetation data (following Bråthen et al. 2007), and average density from 2003 to 2019 as a predictor for the 2020 vegetation data, to not overlap with the previous interval. Results were not sensitive to the length of the intervals (2003-2019 vs. 2000-2019).

### Vegetation sampling

We used the point-intercept method (Bråthen and Hagberg 2004) with a triangular 3-pin frame to obtain a measure of vascular plant abundance. We counted all hits of all vascular species in 11 0.08m ^2^ plots spaced along each 50m transect. Prior to further analysis, we converted point-frequency hits per species per plot to biomass estimates (g/m ^2^) using established calibration equations (Tuomi et al. 2018; Ravolainen et al. 2010). For analyzing biomass data, we first pooled species-specific biomass estimates based on functional grouping to forbs, graminoids, deciduous woody shrubs and dwarf-shrubs (deciduous woody), and crowberry. Other groups not included in the analysis (Supplementary Result Table 1A) were other evergreen woody dwarf-shrubs, non-woody evergreen plants and vascular cryptogams. Other evergreen woody dwarf-shrubs were not included in statistical analyses as responded very similarly to crowberry but have comparably low biomass. We then averaged biomass of all 11 plots along each transect to reach a community-averaged estimate of g/m ^2^ for forbs (non-zero sample size N _year_: N_2003_ = 136, N_2020_ = 136), graminoids (N_2003_ = 257, N _2020_ = 252), deciduous woody (N _2003_ = 282, N _2020_ = 279), and crowberry (N _2003_ = 261, N_2020_ = 269). We also calculated the cover of the functional groups within each community, using plot-level presence-absence data and summarized numbers of plots with each functional group present in each transect.

### Statistical analyses

To test our hypothesis, we fitted Bayesian linear multilevel gamma-hurdle models with the package *brms* (Bürkner 2017) in the R statistical environment (R Core Team 2021) (version 4.0.4/15.02.2021 and later). We decomposed the reindeer density to its spatial, temporal, and residual components (Oedekoven et al. 2017), and standardized all three predictors to a mean of 0 and variance of 1 for better effect comparability and model convergence. The temporal component represents the change in average reindeer densities over two time points, the spatial component the difference associated with the district reindeer densities averaged over time and the residuals the district- and year specific variation in reindeer densities from the spatial and temporal averages. To fully test the expectations in Fig. 1e, models for forbs and graminoids included a quadratic term for the spatial component, while deciduous dwarf-shrubs and crowberry were included linear term only, and all models included block and district as group-level intercepts. We fitted the Bayesian generalized linear mixed models with weakly informative default priors (Bürkner 2017) and checked model convergence and independence of HMC chains based on the 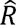 statistics (< 1.002) and effective sample size (> 2000) (Muth, Oravecz, and Gabry 2018). In addition, we fitted negative-binomial hurdle models for plant functional group cover in the same way as described above for biomass data. We used package default weakly informative priors for population and group-level predictors and family parameters (Bürkner 2017). We confirmed model fit visually through posterior predictive checks as well as comparing model-simulated data to observed data (Muth, Oravecz, and Gabry 2018). Lack of spatial autocorrelation in group-level effects and model residuals was assessed visually. Data visualization was done with packages ggplot2 (Wickham 2009) ggdist (Kay 2021) and ggpmisc (Aphalo 2023).

## Author contributions

K.A.B. conceived the idea, M.T., K.A.B. and N.Y. planned the re-sampling design. K.A.B. designed conceptual figures with support from M.T., and based on discussions with all authors. M.T., K.A.B., T.Aa.U., N.Y. and S.Z. collected the data. M.T., K.A.B. and N.Y. planned data analysis. M.T. analyzed the data and extracted the results. M.T. and K.A.B. led data interpretation, with all authors contributing to interpreting the results. M.T. wrote the manuscript with and support from K.A.B. All authors contributed substantially to editing the manuscript.

## Acknowledgements

The work was funded by the Norwegian Research Council (FRIPRO project MONEC, code 302749 to K.A.B.). We thank Nhat Minh Pham for comments on previous drafts of the manuscript, and Karoline Helene Aares, Hanna Böhner, Lea Lipphardt, Hans Ivar Hortmann, Kinga Skalska, Katrine Skamfer Hoset and Sindre Natvik for conducting field work, and the Norwegian Coast Guard, especially the crew of KV Farm, for their hospitality and invaluable logistic help during the field work campaign. We thank Audun Stien for discussions on the calf weight-density relationship. We also wish to thank MONEC partners for discussions on implications of crowberry encroachment for their reindeer and sheep herding SES. M.T. thanks the Mikkeli University Consortium (MUC) for providing working facilities.

