## Supplementary materials for "Greening conceals evergreening: contrasting trends for a socio-ecological system in Arctic Europe"

Supplementary Content Table

Literature synthesis of key characteristics of Empetum nigrum (ssp. hermaphroditum) that underlie the dominance, niche constructing ability and encroachment capacity of the species. For references before 2000, we mainly refer to (Tybirk et al., 2000; Wardle et al., 1998).

| Life history | | |
| --- | --- | --- |
| Growth and dispersal | Dual propagation strategy: infilling and horizontal spread through seeds, monopolizing resources through clonal growth and shoot elongation | (Angers-Blondin & Boudreau, 2017; Szmidt et al., 2002; Wardle et al., 1998) |
|  | Seed dispersal through endozoochory | (Bråthen et al., 2007) |
|  | Potentially very long-lived clonal plant (140 yr) | (Bell & Tallis, 1973) |
|  | Forms dense and extensive mats with horizontal but shallow roots | (Bell & Tallis, 1973; Tybirk et al., 2000) |
|  | High phenotypic plasticity to meet varying habitat (e.g. snow) conditions | (Bienau et al., 2014) |
| Community and ecosystem effects (niche construction and legacy effects) | | |
| Competition | [in nutrient poor and low-pH soils] Competitively superior to herbaceous plants; more neutral or facilitative to plants with high LDM content | (Mod et al., 2014; Pellissier et al., 2010) |
|  | As an ericoidmycorrhizal plant, assimilates N from organic N forms; yet competes effectively for NH_4_-N | (Aerts, 2003, 2010; Tybirk et al., 2000) |
| Palatability | Low palatability of old vegetative tissues, reindeer and other herbivores avoid and have low densities in areas with high *Empetrum* cover | (Ims et al., 2007; Iversen et al., 2014; Tybirk et al., 2000; Wardle et al., 1998) |
|  | Berries and young leaves are eaten by reindeer and other herbivores | (Bråthen et al., 2007) |
| Allelopathy | Phytotoxic phenols (e.g. batatasin III) present in leaves and litter hamper germination, nutrient uptake and growth of other plants and ectomycorrhizal tree seedlings | (Bråthen et al., 2010; González et al., 2015; M. C. Nilsson et al., 2000; M.-C. Nilsson et al., 2011; Tybirk et al., 2000; Wallstedt et al., 2001; Wardle et al., 1998) |
|  | Allelopathic legacy effect of *Empetrum* litter on plant establishment and growth can last at least for 10 years, even if above-ground plant is removed | (Aerts, 2010; González et al., 2021; Wardle & Jonsson, 2014) |
|  | Alters succession and reduces forest productivity | (Tybirk et al., 2000; Wardle et al., 1998) |
| Effects on floral biodiversity | Suppression of diversity of herbaceous plants | (Bråthen & Ravolainen, 2015; Mod et al., 2014; Pellissier et al., 2010) |
|  | “Gatekeeper” when present as >¼ of standing vascular crop – curbs warming-induced increases in tundra species richness | (Bråthen et al., 2018) |
| Effect on soil nutrients, microbes and processes | Negative effects on soil nematodes, microbial biomass, nutrient mineralization an decomposition rates | (Castells et al., 2005; M.-C. Nilsson & Wardle, 2005; Ruess et al., 1998; Tybirk et al., 2000; Wardle et al., 1998) |
|  | High phenolic litter, promotes humus where N locked in organic forms | (Tybirk et al., 2000; Wardle et al., 1998) |
| Responses to the biotic and abiotic environment | | |
| Response to disturbance and herbivory | Herbivore trampling, small-rodent activity and anthropogenic disturbance (e.g. ATV’s) cause mortality | (Angers-Blondin & Boudreau, 2017; Bell & Tallis, 1973; Olofsson, 2009; Olofsson et al., 2012; Tuomi et al., 2018; Väisänen et al., 2014) |
|  | Fire often kills *Empetrum*, and in post-fire succession Empetrum becomes dominant after a century or more | (Tybirk et al., 2000; Wardle et al., 1998) |
|  | May be promoted by *Rangifer* through apparent competition and endozoochory at low-intermediate coverage | (Bråthen et al., 2007, 2018) |
|  | Increased shoot mortaility after autumnal moth outbreaks | (Bokhorst et al., 2015) |
|  | Fire and animal defecation can alleviate allelopathic legacy effects of Empetrum litter on germination and growth of herbaceous plants [possibly linked with increased soil pH] | (Bråthen et al., 2010; Keech et al., 2005; Wardle et al., 1998) |
| Response to warming | Extreme winter warming events can reduce shoot growth and cause mortality | (Bokhorst et al., 2015, 2011) |
|  | Vulnerability to specialist fungus *Arwidssonia empetri* under increased snow cover | (Olofsson et al., 2011) |
|  | Susceptible to drought at southern distribution margin | (Hein et al., 2021) |
|  | Tolerates well ice encasement | (Preece et al., 2012; Preece & Phoenix, 2014) |
|  | Sexual reproduction and colonization promoted in a warming climate | (Angers-Blondin & Boudreau, 2017) |
|  | Warming leads to positive growth response and shoot elongation | (Buizer et al., 2012; Kaarlejärvi et al., 2012; Klanderud & Birks, 2003; Wada et al., 2002; Wilson & Nilsson, 2009) |
| Response to nutrient addition | Increased abundance with single addition of inorganic N and enhanced recovery from mechanical disturbance | (Aerts, 2010; JONASSON, 1992) |
|  | Repeated inorganic N addition leads to (resident) graminoids out-competing *Empetrum* | (Eskelinen et al., 2012; M.-C. Nilsson et al., 2002) |
|  | Repeated P (N+P) addition can promote *Empetrum* | (Wardle et al., 2016) |

Supplementary Results Tables

Supplementary Table 1.

**A)** Average biomass (± 95% confidence intervals) and species richness of all plant forage groups, calculated from raw observed values. **B)** Average cover (± 95% confidence intervals, as number of plots present per transect) of studied plant forage groups and other evergreen dwarf-shrubs, calculated from raw observed values.

| A |  |  |  |  |
| --- | --- | --- | --- | --- |
| Functional group | Biomass ± CI -03 | Biomass ± CI -20 | Richness -03 | Richness -20 |
| Deciduous woody | 83.927 ± 9.228 | 111.987 ± 11.737 | 12 | 12 |
| Crowberry | 141.265 ± 13.839 | 235.691±21.979 | 1 | 1 |
| Other evergreen woody | 28.723 ± 4.683 | 45.246 ± 8.062 | 10 | 10 |
| Evergreen non-woody | 0.517 ± 0.251 | 0.612 ± 0.492 | 6 | 5 |
| Forb | 4.658 ± 1.113 | 4.472 ± 1.533 | 43 | 46 |
| Graminoid | 24.982 ± 3.75 | 22.854 ± 3.488 | 29 | 35 |
| Vascular cryptogam | 0.581 ± 0.320 | 0.418 ± 0.196 | 6 | 8 |
| B |  |  |  |  |
| Functional group | Cover ± CI -03 | Cover ± CI -20 |  |  |
| Deciduous woody | 4.873 ± 0.303 | 4.884 ± 0.312 |  |  |
| Crowberry | 4.699 ± 0.351 | 5.521 ± 0.349 |  |  |
| Other evergreen woody | 1.901 ± 0.232 | 2.25 ± 0.24 |  |  |
| Forb | 1.264 ± 0.230 | 1.182 ± 0.211 |  |  |
| Graminoid | 4.065 ± 0.352 | 3.798 ± 0.344 |  |  |

Supplementary Table 2.

Model parameter estimates and credible intervals, with plant functional group cover as response and decomposed reindeer density as predictors. Note that the temporal coefficient links with decrease in reindeer density over time, and hence a negative value indicates an increase over time. Error terms Q2.5 and Q97.5 represent the 95% credible interval. Bold font indicates strong support for the effect, and italic indicates relatively strong support for the effect, given the data.

| **Response** | **Parameter** | **Estimate** | **Q2.5** | **Q97.5** |
| --- | --- | --- | --- | --- |
| Cover ~ spatial + spatial^2^ + temporal + residual | | | | |
| Forbs |  |  |  |  |
|  | Intercept | 0.303 | -0.159 | 0.748 |
|  | Spatial | -0.423 | -0.913 | 0.045 |
|  | Spatial^2^ | 0.135 | -0.256 | 0.453 |
|  | Temporal | 0.036 | -0.130 | 0.205 |
|  | Residual | -0.070 | -0.187 | 0.045 |
| Graminoids |  |  |  |  |
|  | Intercept | 1.370 | 1.153 | 1.587 |
|  | Spatial | -0.035 | -0.278 | 0.202 |
|  | Spatial^2^ | 0.037 | -0.122 | 0.192 |
|  | Temporal | 0.041 | -0.034 | 0.116 |
|  | Residual | -0.018 | -0.067 | 0.031 |
| Cover ~ spatial + temporal + residual | | | | |
| Deciduous woody |  |  |  |  |
|  | Intercept | 1.569 | 1.435 | 1.703 |
|  | Spatial | 0.036 | -0.108 | 0.185 |
|  | Temporal | -0.012 | -0.066 | 0.042 |
|  | Residual | -0.019 | -0.057 | 0.017 |
| Crowberry |  |  |  |  |
|  | Intercept | 1.661 | 1.546 | 1.769 |
|  | Spatial | 0.062 | -0.066 | 0.189 |
|  | **Temporal** | **-0.103** | **-0.157** | **-0.047** |
|  | Residual | 0.000 | -0.037 | 0.037 |

Supplementary Table 3.

Estimated standard deviations from the population-level estimate of group-level (random) intercepts of the hierarchical linear models. Error terms Q2.5 and Q97.5 represent the 95% credible interval, within which the true estimate lies with a 95% probability. A) Model with plant group biomass as the response. B) Model with plant group cover as the response.

| Response | Parameter | Estimate | Q2.5 | Q97.5 |
| --- | --- | --- | --- | --- |
| A) Biomass | | | | |
| Forbs | Block (intercept) | 0.284 | 0.074 | 0.501 |
|  | District (intercept) | 0.545 | 0.267 | 0.887 |
| Graminoids | Block (intercept) | 0.558 | 0.392 | 0.759 |
|  | District (intercept) | 0.374 | 0.046 | 0.738 |
| Dedicuous woody | Block (intercept) | 0.358 | 0.238 | 0.507 |
|  | District (intercept) | 0.441 | 0.258 | 0.698 |
| Crowberry | Block (intercept) | 0.423 | 0.323 | 0.542 |
|  | District (intercept) | 0.117 | 0.004 | 0.314 |
| B) Cover | | | | |
| Forbs | Block (intercept) | 0.196 | 0.012 | 0.458 |
|  | District (intercept) | 0.621 | 0.296 | 1.082 |
| Graminoids | Block (intercept) | 0.344 | 0.229 | 0.483 |
|  | District (intercept) | 0.244 | 0.037 | 0.463 |
| Dedicuous woody | Block (intercept) | 0.220 | 0.140 | 0.321 |
|  | District (intercept) | 0.230 | 0.121 | 0.367 |
| Crowberry | Block (intercept) | 0.253 | 0.175 | 0.345 |
|  | District (intercept) | 0.134 | 0.009 | 0.285 |

Supplementary Figures

Supplementary Figure 1.

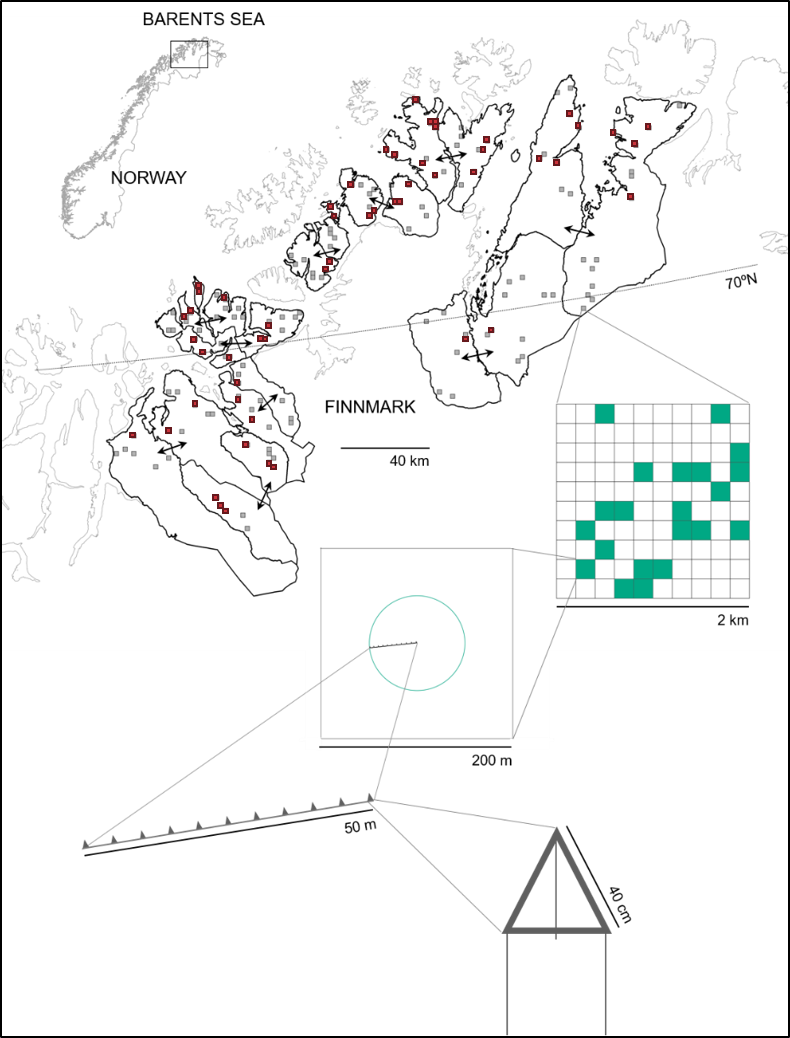

**Supplementary Figure 1.** The study and sampling design as described during the first survey (Bråthen et al 2007). The areas sounded with black lines are reindeer herding summer districts. Red small squares indicate 2x2km landscape areas sampled both in 2003 and 2020, grey small squares the original survey.

Supplementary Figure 2.

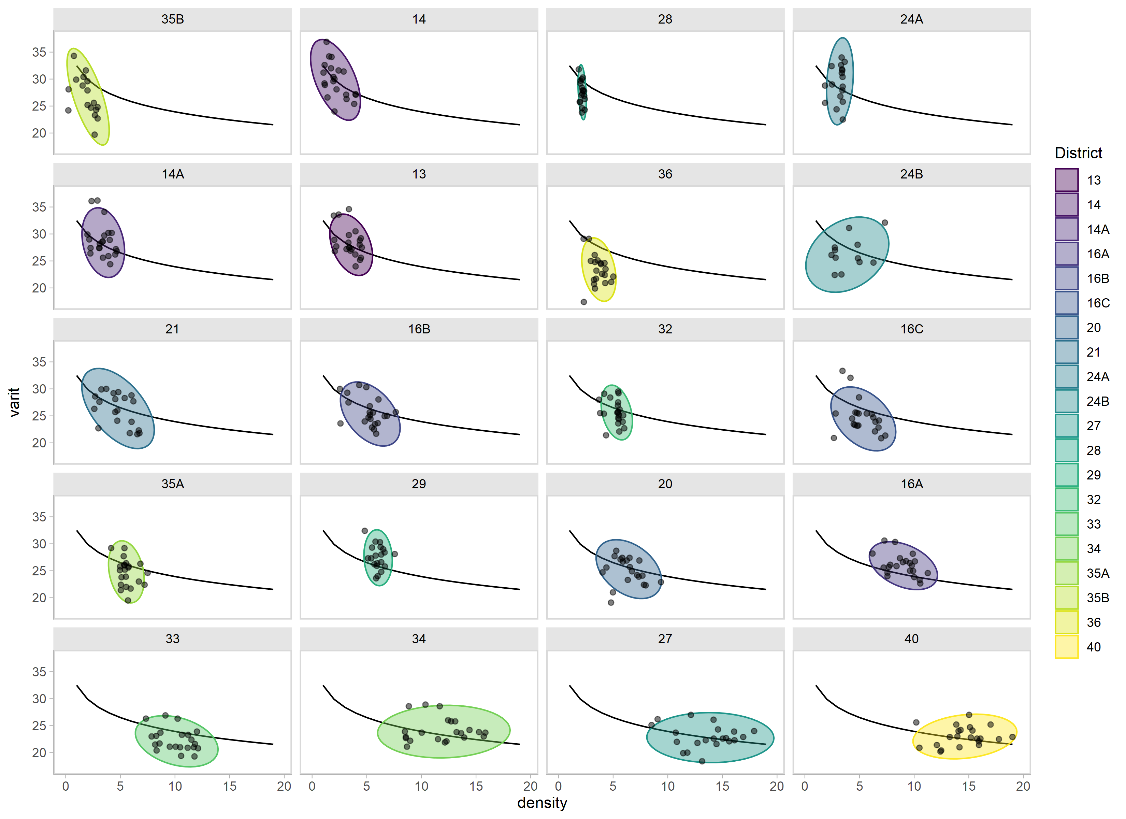

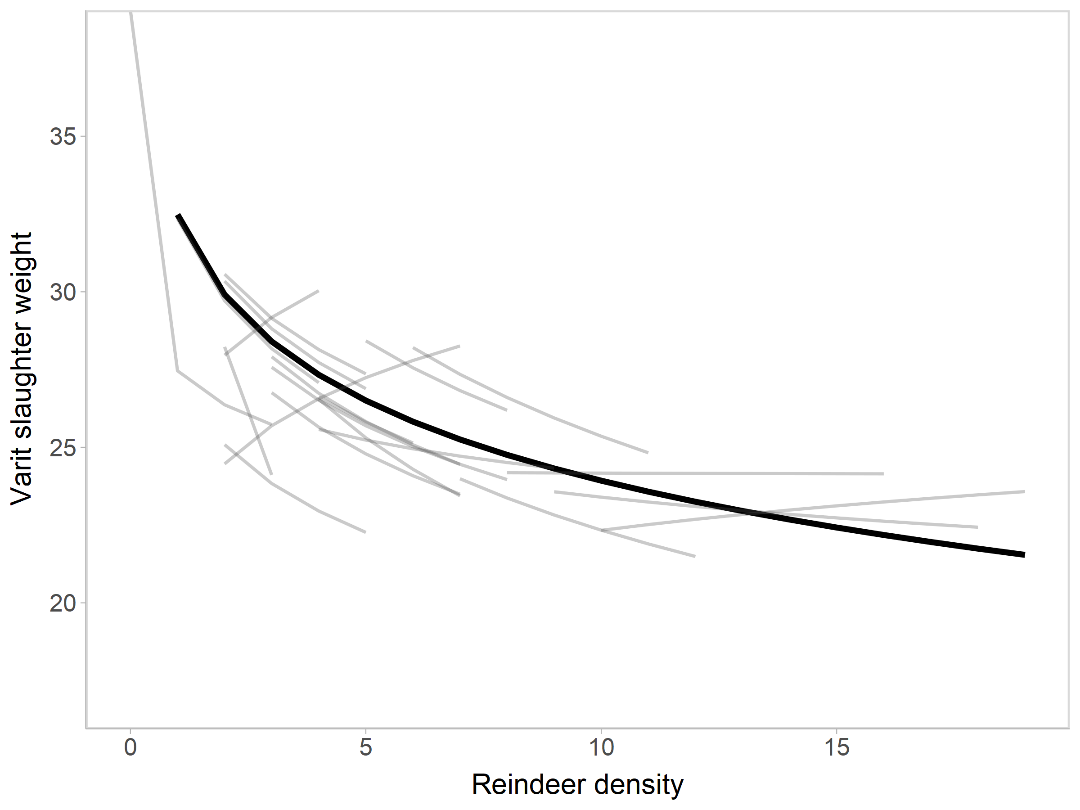

Varit slaughter weight (kg)

Reindeer density (animals/km^2^)

Reindeer density (animals/km^2^)

Varit slaughter weight (kg)

**a**

**b**

**Supplementary Figure 2** Relationship between reindeer density and varit (1.5 years old calves) body mass in the studied reindeer districts during the study period (Ministry of Agriculture, via reinbase.no). **A)** Similar to calf weights, varit weights are also used as an indicator in the management system (Reindeer Herding Act; Veileder for fastsetting av økologisk bærekraftig reintall; White Paper 32(2016-2017)). Lognormal relationship across all districts (black line) indicates a negative density dependence of varit weights, while data within each district (grey lines) indicates variability from this general relationship. **B)** Observations (dark points) in each district plotted with the lognormal regression line across districts (black line), with normal data ellipses indicating distribution of observations within each district.

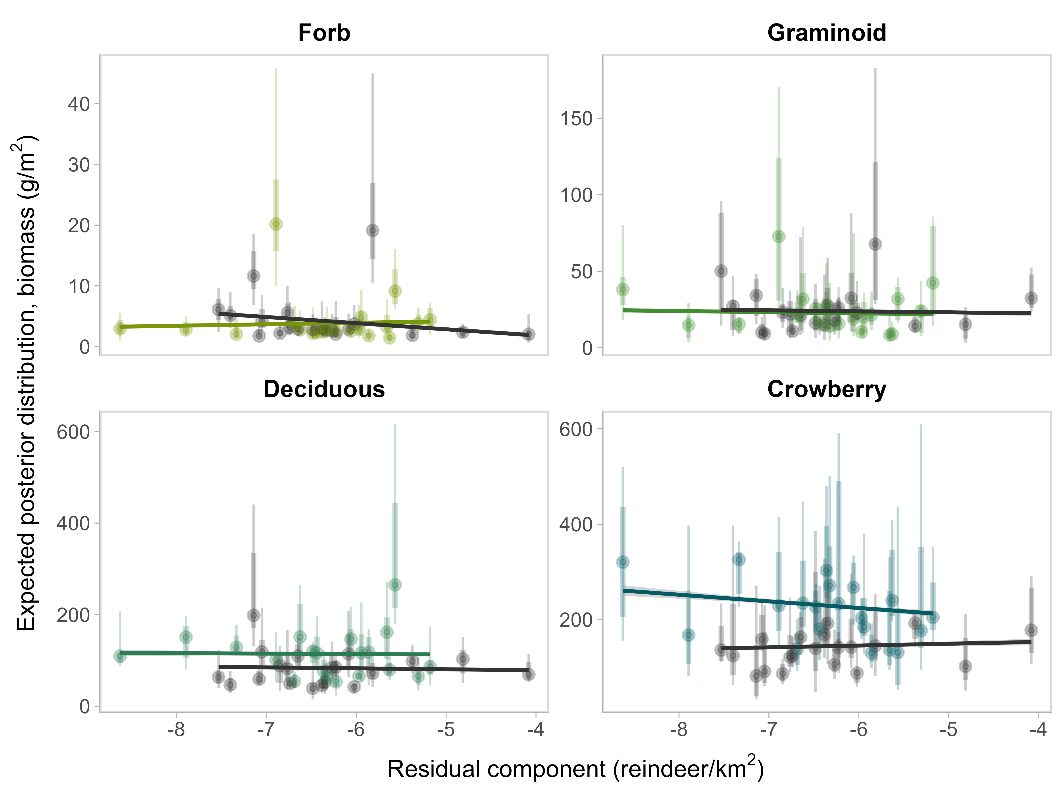
Supplementary Figure 3

**Supplementary Figure 3.** Association (mean and model-derived 95% and 80% credible intervals of the estimated posterior mean distribution) between plant groups and the district-specitic changes in reindeer density (the residual component). Each point represents a district and year, with greens representing districts in 2020 and grey in 2003. The coloured regression lines across all districts are estimated with the package ggpmisc function stat_poly_line (Aphalo 2023)

Supplementary Figure 4.

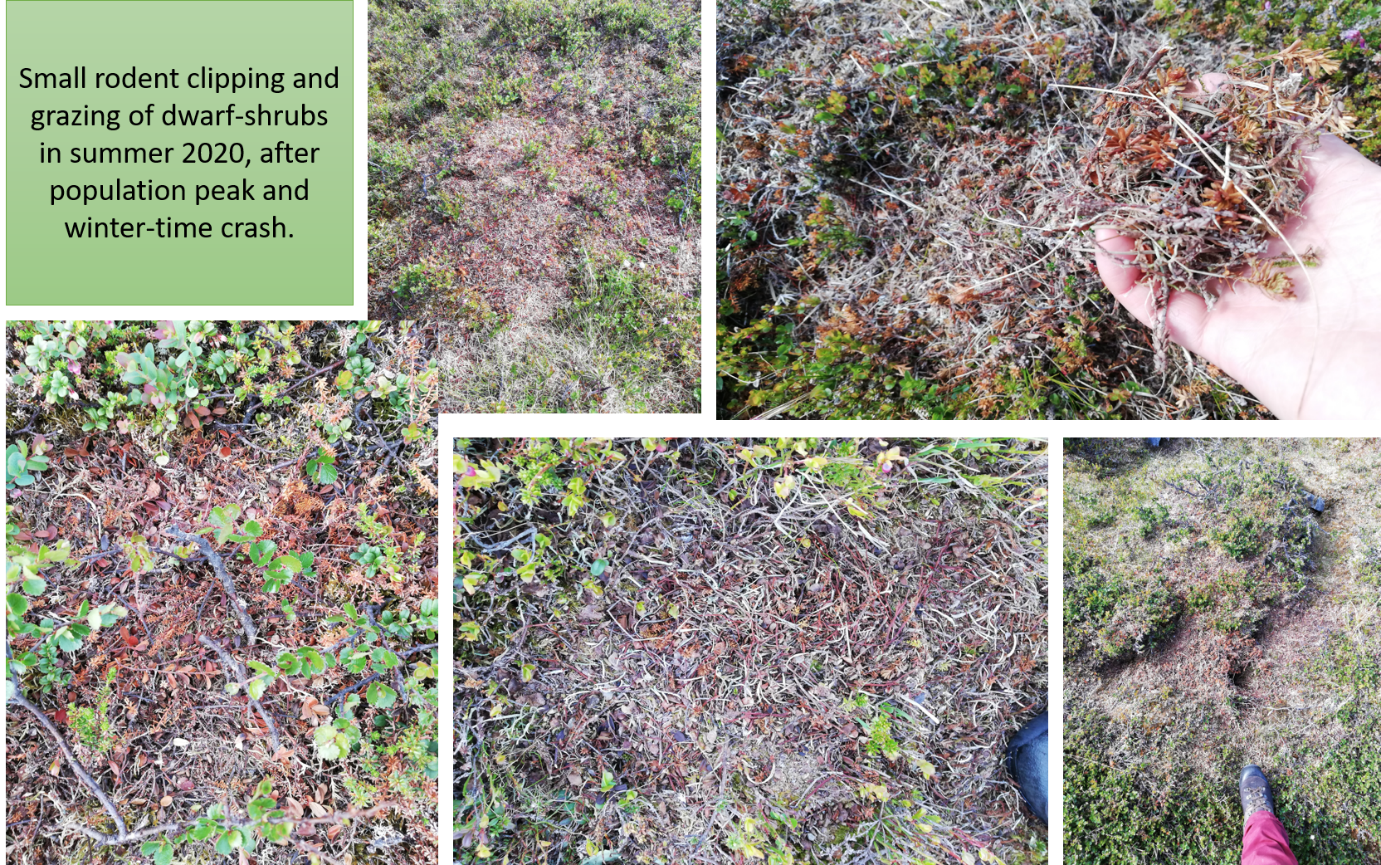

**Supplementary Figure 4.** Small rodent grazing marks and damage in dwarf-shrub vegetation after population peak winter. Small rodent peak and crash during winter 2019-2020 left visible clipping and grazing marks in the tundra. Especially dwarf-shrubs Vaccinium myrtillus, Empetrum nigrum and Betula nana were visibly damaged.
